# Impact of > 24 h sustained wakefulness and subsequent recovery sleep on time-dependent changes in microRNA factors of individuals with post-acute phase mild traumatic brain injury

**DOI:** 10.1101/2025.09.11.675463

**Authors:** Allison Brager, Katie Edwards, Cassie Pattinson, J. Peyer, Jessica Gill

## Abstract

**Introduction:** The purpose of the present study was to determine whether mild traumatic brain injury (mTBI+) results in latent changes in microRNA expression profiles across an episode of total sleep deprivation (TSD).

**Methods:** Seven previously concussed (mTBI+) adults (24.5 ± 5.3 y.o.) and 6 non-concussed control adults (mTBI−; 24.8 ± 1.6 y.o.) underwent 24 h TSD (T2) preceded by 8 h baseline sleep (T1; BSL) and followed by 8 h recovery (T3; REC) sleep. Salivary microRNA expression was assessed across the entire study.

**Results:** Subjects (mTBI+ and mTBI−) had differential expression of salivary microRNA targets across TSD. mTBI+ subjects had greater change to microRNA expression profiles compared to mTBI− subjects between T1 and T3.

**Discussion:** Although there is some evidence that TSD may unmask latent changes in gene expression in mTBI+ subjects, a definitive conclusion was precluded by differences in baseline sleep in mTBI+ (vs. mTBI−) subjects measured through polysomnography (not shown). However, this study is unique in that the mTBI+ subjects were exposed to a sleep deprivation challenge very shortly after medical clearance from mTBI demonstrating lingering neurobiological impacts of mTBI.

## Introduction

Sleep complaints are persistent and predictive of long-term recovery post-acute phase mild traumatic brain injury (mTBI). Sleep disturbances are also prevalent among military personnel, especially individuals with mild traumatic brain injury (mTBI).^1,2^ While much is known about the impact of mTBI on sleep architecture, less is known about the impact of mTBI on sleep homeostatic and circadian processes, including downstream molecular cascades. To better characterize these effects, we determined and isolated mTBI effects on microRNA expression across baseline sleep, a sleep deprivation challenge, and subsequent recovery. microRNAs (miRNAs) are small (~22 nucleotide) non-coding RNA strands that base pair with messenger RNA (mRNA) to degrade or inhibit its translation into protein. Sleep loss has previously been shown to induce changes to miRNA^3,4^.

## Methods

### Participants

Study participants underwent an intensive pre-screen at Walter Reed Army Institute of Research (WRAIR) to identify healthy sleepers. All study participants were between the ages of 18-39 (n = 17 enrolled; n = 13 completed study; 8 females; mean age, 26.0 ± 1.5). mTBI+ participants (n = 7; mean age, 26.1 ± 2.5) had been diagnosed with mTBI within 3-12 months prior to enrollment. Controls (n = 6; mean age, 24.8 ± 1.6) had no history of mTBI for a minimum of 5 years prior to enrollment.

### Concussion History and Symptomatology

Sports-related trauma (57.1%) was the primary cause of mTBI in this study. Secondary causes included blunt trauma (14.3%), motor vehicle crash (14.3%), and fall (14.3%). The Rivermead questionnaire (King et al., 1995) - the gold standard self-report of concussion symptomatology (scale of 0-4; 0 = Not experienced at all; 4 = A severe problem) - was administered during screening to all participants. Average Rivermead scores during screening (baseline) was 2.8 ± 0.8 for all participants (mTBI+ or mTBI−; n = 13) and did not differ between study groups (t-test: p > 0.05).

### Experimental Timeline

For two weeks prior to entering the sleep laboratory, wrist actigraphy (Phillips Respironics; Murrysville, PA; not shown) was used to verify that habitual sleep durations and timing were within normal limits (7-9 h time in bed). When in the sleep laboratory, all scheduled sleep was polysomnographically recorded with 8-lead (gold cup) electrodes affixed to the scalp using the 10-20 system, chin (to measure EMG), and the outer canthi of the eyes (to measure eye movements; not shown). Following one overnight sleep opportunity with 8 h time in bed (T1; baseline; 1100 - 0700 EST), wakefulness was maintained continuously for 40 h (T2). Participants were then allowed an 8 h sleep opportunity (T3; recovery sleep; 1100 – 0700 EST). Every 4 hours of the study, saliva was collected for microRNA analysis.

### Micro-RNA Sequencing and Analysis

Saliva samples were collected and immediately frozen every 4 h at the start of the TSD period (T1) and continued until the end of the TSD period (T2) and again following the 8 h recovery sleep period (T3). miRNA expression was determined with the nCounter® Human v3 miRNA Expression Panels (NanoString Technologies, WA, USA), each of which consists of 800 miRNA probes. After code count normalization of nCounter® Human v3 miRNA Expression Panels, miRNAs with an adjusted Bonferroni p-value <0.05 were considered as differentially expressed.

### Statistics

All statistical analyses for neurobehavioral and sleep physiological outputs were performed using SPSS v26 (Chicago, IL). A p-value < 0.05 with adjusted Bonferroni comparisons was deemed statistically significant. The types of statistical analysis used are specified in the results section.

## Results

### MicroRNA expression was altered throughout the course of total sleep deprivation (TSD) in concussed subjects only

A summary of between-subject (a) and within-subject timepoint interrelationships (b – controls, mTBI−; c – concussed subjects, mTBI+) can be found in **Table 1**. MicroRNA expression was assessed at three time points: T1 (start of TSD); T2 (end of TSD); and T3 (end of recovery sleep). There were no changes to microRNA expression targets at T2 (vs. T1; end of TSD) for mTBI− subjects (p> 0.05). In mTBI+ subjects, 4 microRNA targets (see **Table 1**) were altered in expression at T2 (vs. T1; end of TSD; p <0.05, all). Database queries (NCBI Gene) demonstrated that most of the affected microRNAs play a role in controlling cell growth/proliferation and apoptosis, with many linked to various cancers (acting as either tumor suppressors or oncogenes depending on the specific cancer type and context), and some known to be multifunctional (i.e., implicated in diverse biological processes including the immune response, inflammation, and fibrosis; Irwin et al., 2016).

**Table 1.**
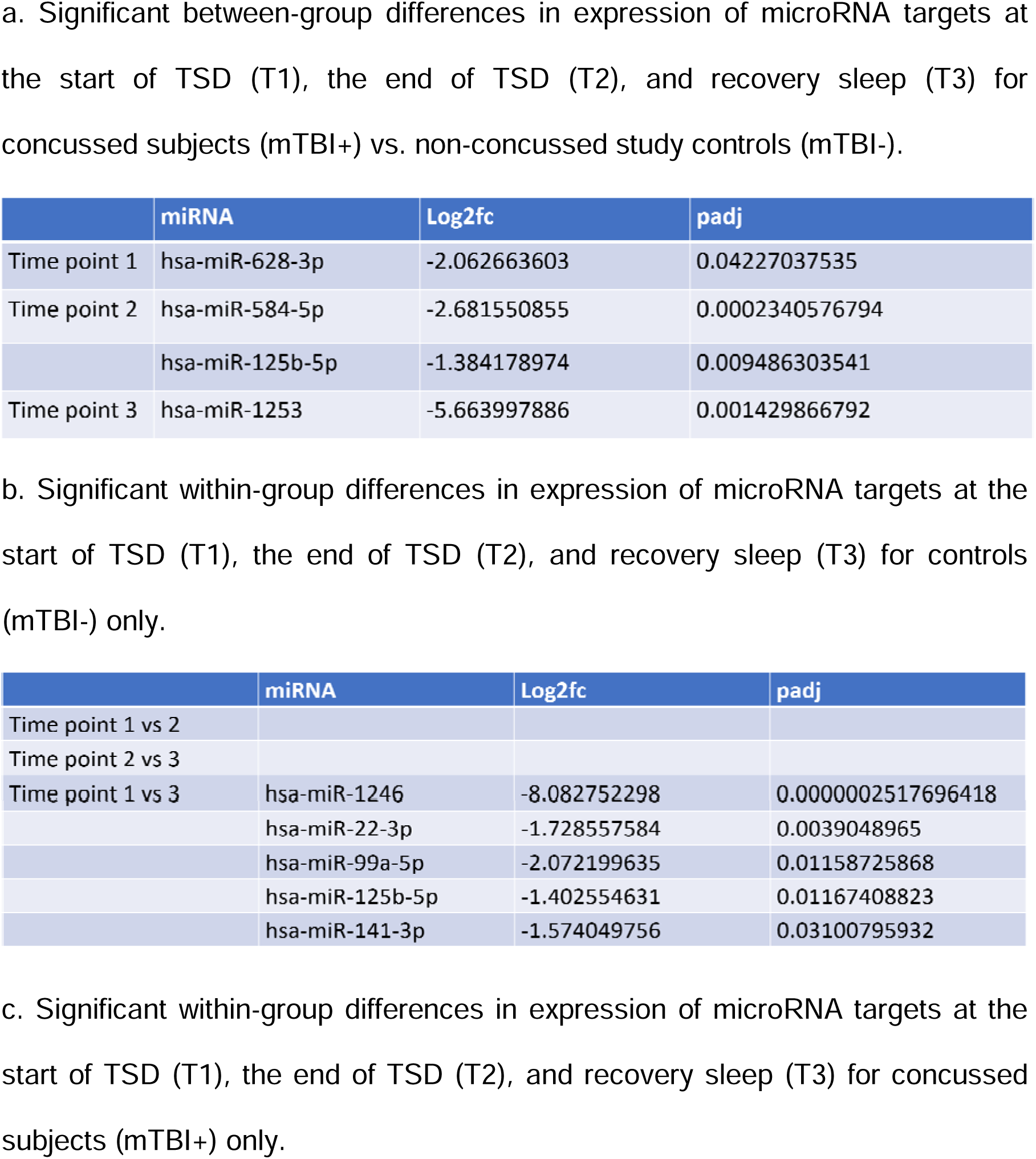
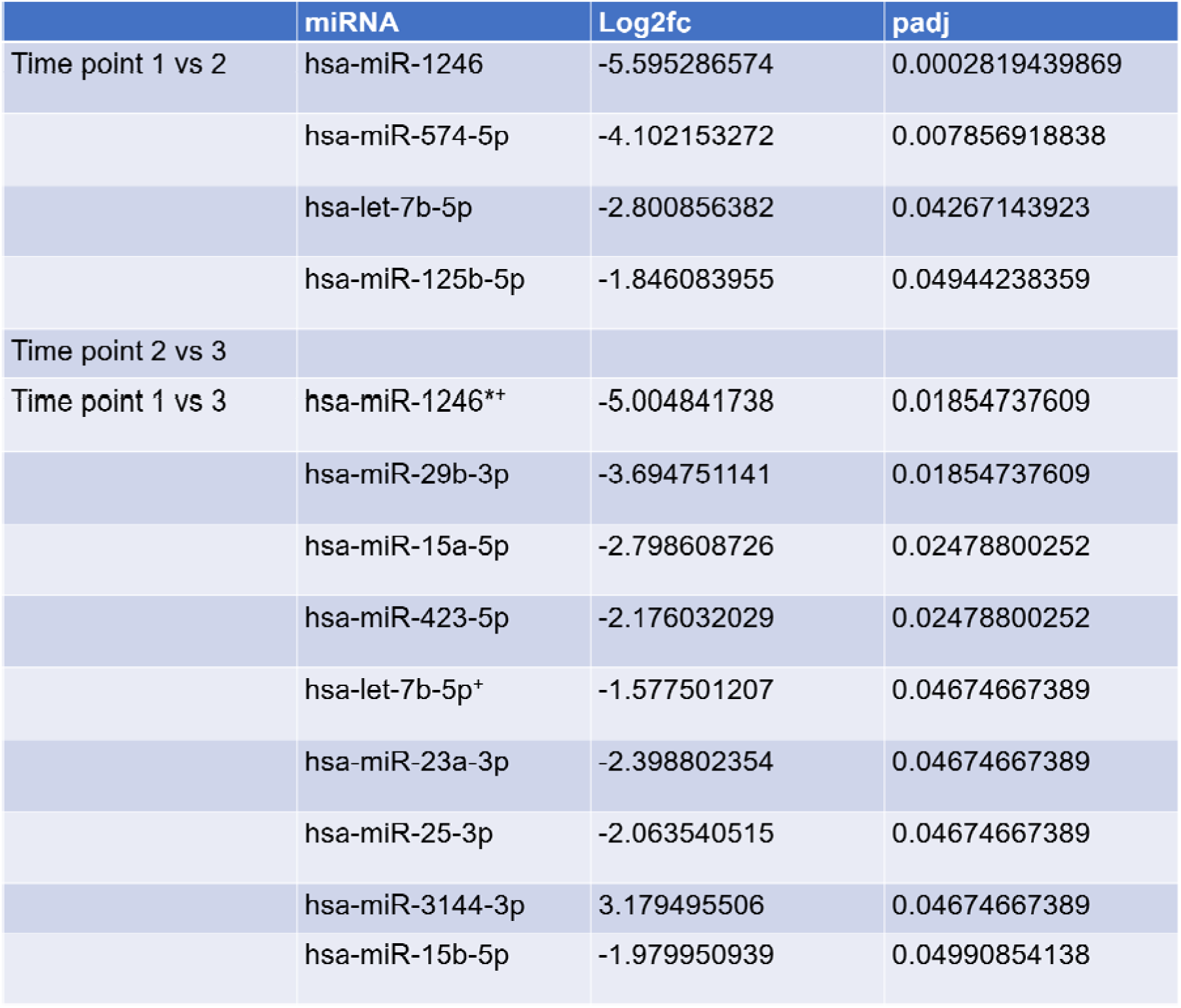
A summary of salivary microRNA targets showing between-subject (a) and within-subject comparisons (b – mTBI−; c – mTBI+) across T1 (start of TSD), T2 (end of TSD), and T3 (end of recovery sleep).

## Discussion

Aligned with other studies showing altered microRNA expression in humans subjected to acute sleep deprivation (Irwin et al., 2016; Gaine et al., 2018), our results also show that TSD unmasks differences in molecular factors in mTBI+ subjects that were not prevalent in mTBI− controls (**Table 1**). These data present a first-ever look at the combined effects of post-acute phase mTBI and TSD on miRNA-regulated gene expression in humans to the best of our knowledge. Continued research is needed to elucidate biomarkers and subsequent underlying biological mechanisms which are involved in sleep processes during the post-acute phase of mTBI.

To conclude, although there is some evidence that TSD may unmask latent changes in gene expression in mTBI+ subjects post-acute phase of mTBI, a definitive conclusion was precluded by differences in baseline sleep in mTBI+ (vs. mTBI−) controls (data not shown), suggesting that mTBI+ subjects may habitually carry a relatively elevated sleep debt (vs. mTBI− controls). Future studies should include a higher saliva sampling rate to better characterize the differential effects of sleep loss on mRNA expression in mTBI+ vs. mTBI− subjects. Overall, an improved understanding of sleep-wake/circadian dynamics during the post-acute phase of mTBI can help with the integration of prophylactic and therapeutic treatments post-acute phase mTBI, particularly for military personnel and related communities who do not have luxuries of regular and sufficient sleep.

## Acknowledgments

The authors would like to thank every member of the WRAIR Sleep Research Center “Dream Team;” clinical research coordinators, lab technicians, registered PSGTs, and human subjects research support staff. None of this work could be completed without their dedication and selfless service.

Material has been reviewed by the Walter Reed Army Institute of Research. There is no objection to its presentation and/or publication. The opinions or assertions contained herein are the private views of the author, and are not to be construed as official, or as reflecting true views of the Department of the Army or the Department of Defense.

## References

1. Mysliwiec V, Gill J, Lee H, Baxter T, Pierce R, Barr TL, et al. Sleep disorders in US military personnel: A high rate of comorbid insomnia and obstructive sleep apnea. Chest. 2013;144(2):549–57.

2. Capaldi V, Guerrero M, Killgore W. Sleep Disruptions among Returning Combat Veterans from Iraq and Afghanistan. Mil Med. 2010;176(August 2011):879–88.

3. Gaine M, Chatterjee S, Abel T. Sleep Deprivation and the Epigenome. Front Neural Circuits. 2018; 12:14.4.

4. Irwin M, Olmstead R, Carroll J. Sleep Disturbance, Sleep Duration, and Inflammation: A Systemic Review and Meta-Analysis of Cohort Studies and Experimental Sleep Deprivation. Biol Psychiatry. 2016 Jul 1; 80(1): 40–52.

